# Salience Network Disruption in US Army Soldiers with PTSD

**DOI:** 10.1101/496505

**Authors:** Chadi G. Abdallah, Christopher L. Averill, Amy E. Ramage, Lynnette A. Averill, Selin Goktas, John H. Krystal, John D. Roache, Patricia Resick, Stacey Young-McCaughan, Alan L. Peterson, Peter Fox, the STRONG STAR Consortium

## Abstract

**BACKGROUND:** Better understanding of the neurobiology of posttraumatic stress disorder (PTSD) may be critical to developing novel, effective therapeutics. Here, we conducted a data-driven investigation using a well-established, graph-based topological measure of nodal strength to determine the extent of functional dysconnectivity in a cohort of active duty US Army soldiers with PTSD compared to controls.

**METHODS:** 102 participants with (n=50) or without PTSD (n=52) completed functional magnetic resonance imaging (*f*MRI) at rest and during symptom provocation using subject-specific script imagery. Vertex/voxel global brain connectivity with global signal regression (GBCr), a measure of nodal strength, was calculated as the average of its functional connectivity with all other vertices/voxels in the brain gray matter.

**RESULTS:** In contrast to during resting-state, where there were no group differences, we found a significantly higher GBCr, in PTSD participants compared to controls, in areas within the right hemisphere, including anterior insula, caudal-ventrolateral prefrontal, and rostral-ventrolateral parietal cortices. Overall, these clusters overlapped with the ventral and dorsal salience networks. *Post hoc* analysis showed increased GBCr in these salience clusters during symptom provocation compared to resting-state. In addition, resting-state GBCr in the salience clusters predicted GBCr during symptom provocation in PTSD participants but not in controls.

**CONCLUSION:** In PTSD, increased connectivity within the salience network has been previously hypothesized, based primarily on seed-based connectivity findings. The current results strongly support this hypothesis using whole-brain network measure in a fully data-driven approach. It remains to be seen in future studies whether these identified salience disturbances would normalize following treatment.

## INTRODUCTION

Posttraumatic stress disorder (PTSD) is a disabling mental illness with poorly understood pathophysiology and a reported crisis in drug development (1,2). Better understanding of the neurobiology of PTSD may be critical to the development of novel effective therapeutics. To date, early neuroimaging work has identified a number of neural circuit abnormalities in PTSD, and more recently, brain-wide functional connectivity network disturbances have been described (3,4). Building on prior work, the aim of the current report is to identify robust brain biomarkers of PTSD and to gain insight into the PTSD brain network abnormalities at rest and during symptom provocation. To achieve this aim, we conducted a data-driven investigation using a well-established, graph-based, topological measure of nodal strength to determine the extent of functional dysconnectivity in a cohort of active duty US Army soldiers with PTSD compared to controls.

Functional connectivity magnetic resonance imaging (*fc*MRI) is a powerful neuroimaging tool that has been extensively used over the past 2 decades to investigate the role of the brain intrinsic connectivity networks (ICNs) (5). It has been successfully employed both to map the architecture of brain systems in healthy individuals (6–8) and to identify circuits and network disturbances in neuropsychiatric disorders (9). Early studies have mostly focused on hypothesis-driven, seed-based analyses and ICNs during resting-state. More recently, data-driven, graph-based *fc*MRI topological measures have been established and *fc*MRI approaches have been increasingly employed during various task and arousal states (10,11). One essential topological measure that has been widely employed to investigate ICN in health and disease is nodal strength, i.e., a measure reflecting the total amount of connectivity between a node (voxel/vertex or brain region) and the rest of the network (e.g., whole brain) (12). In healthy populations, measures of nodal strength, also known as functional connectivity strength, were found to identify major brain networks (13), to predict cognitive functioning and intelligence (14), to correlate with regional brain activity (15,16), and to be directly linked to glutamate neurotransmission (17).

Global brain connectivity with global signal regression (GBCr) is a robust, well-established measure of nodal strength. GBCr of a voxel is the average of its functional connectivity with all other voxels in the brain gray matter. During resting-state, reduced prefrontal GBCr and other comparable measures have been reported in depression (17–21) and in several psychiatric disorders with a considerable chronic stress component (22–25). Moreover, the stress-related prefrontal GBCr deficits were found to normalize following ketamine treatment (17,18,26). Together, these findings have led to the hypothesis that the identified prefrontal GBCr abnormalities may reflect, at least partially, an underlying stress-related synaptic loss and dysconnectivity that have long been reported in preclinical studies of trauma and chronic stress (1,27). Surprisingly, we previously found no prefrontal resting-state GBCr abnormalities in US military veterans suffering from severe PTSD symptoms (28). However, follow-up exploratory analyses revealed a pattern of reduced prefrontal GBCr in veterans who reported high symptoms of avoidance and numbing over the past month but increased prefrontal GBCr in those who reported high arousal symptoms (28). Thus, we speculated that the trauma- and stress-related prefrontal GBCr dysconnectivity may have been masked by arousal-induced increases in cortical GBCr. Consistent with this hypothesis, GBCr is known to be directly affected by acute brain functions as evident by (1) an increased GBCr during treatment with ketamine (a drug known to induce transient glutamate neurotransmission), (2) a reduced GBCr by the glutamate release inhibitor lamotrigine, and (3) an association between GBCr and regional brain activity (16,17,24,26,29,30). However, whether provocation of trauma-related symptoms would increase GBCr is not yet known.

In the current report, we aimed to demonstrate this working model by investigating GBCr at rest and during symptom provocation using personalized script imagery in a cohort of individuals with PTSD and a sex-/age-matched non-PTSD comparison group. We first conducted data-driven whole-brain analyses to determine the presence and location of GBCr disturbances in the PTSD group at rest and during symptom provocation. Then, to facilitate the interpretation of the whole-brain findings, we extracted the GBCr values from the identified regions and conducted *post hoc* analyses to determine the effects of task and subgroups. We predicted no prefrontal abnormalities at rest but increased GBCr during symptom provocation.

## METHODS AND MATERIALS

All behavioral and imaging data were acquired from the STRONG STAR data repository (https://tango.uthscsa.edu/strongstar/subs/rpinfo.asp?prj=12). The imaging data and analyses during symptom provocation are new and have not been used in previous reports. The resting state investigation is a novel analysis of a data set that was separately investigated using an Independent Component Analysis (ICA) approach (Vanasse et al., unpublished data, November 2018). Yet, there is no overlap between the GBCr measures investigated in this study and the resting state ICA report.

### Participants

A total of 102 participants with successful scans were investigated (Table 1). Study procedures were approved by institutional review boards and informed consents were completed prior to participation. All participants had no magnetic resonance contraindication and had a negative drug screen on the day of the scan. The PTSD patients met the following criteria: (1) were active duty US Army soldiers, following deployment to or near Iraq or Afghanistan; (2) were 18 years or older; (3) had PTSD diagnosis, as confirmed by a structured interview based on the *Diagnostic and Statistical Manual for Mental Disorder, Fourth Edition* (*DSM-IV*); (4) experienced a Criterion A traumatic event during deployment; (5) if on psychotropic medications, were on stable doses for at least 6 weeks; (6) did not have imminent suicide or homicide risk; (7) did not have psychosis; (8) did not have moderate or severe traumatic brain injury. The control participants were either healthy control (HC) civilians with no Criterion A trauma, or combat control (CC), who were active duty US Army soldiers and met Criterion A during deployment but did not have PTSD. Baseline symptom severity was determined using the PTSD Check List (PCL) for *DSM-IV*, the Beck Depression Inventory (BDI), and the Beck Anxiety Inventory (BAI).

**Table 1.** Demographics and Clinical Characteristics.

|  | PTSD<br>(n = 50) | Combat Control<br>(n = 29) | Healthy Control<br>(n = 23) |
| --- | --- | --- | --- |
|  | Mean (SEM) or N (%) | Mean (SEM) or N (%) | Mean (SEM) or N (%) |
| Age | 32.8 (1.1) | 31.8 (1.1) | 32.4 (2.1) |
| BMI <sup>a</sup> | 28.9 (0.7) | 28.3 (0.6) | 25.8 (0.7) |
| IQ <sup>a</sup> | 98 (1.5) | 99 (2.1) | 110 (2.6) |
| Sex (Male) | 46 (92%) | 27 (91%) | 20 (87%) |
| Race (White) | 32 (64%) | 18 (62%) | 16 (70%) |
| Race (Black) | 11 (22%) | 6 (21%) | 4 (17%) |
| Ethnicity<br>(Hispanic) <sup>a</sup> | 10 (20%) | 10 (35%) | 10 (44%) |
| PCL <sup>a</sup> | 56.7 (1.8) | 20.0 (0.7) | 19.8 (0.8) |
| BDI <sup>a</sup> | 28.2 (1.7) | 2.3 (0.7) | 1.7 (0.5) |
| BAI <sup>a</sup> | 19.3 (1.8) | 1.7 (0.4) | 1.9 (0.4) |

**Abbreviations:** BAI, Beck Anxiety Inventory; BDI, Beck Depression Inventory; BMI, body mass index; IQ, intelligence quotient; PCL, PTSD Checklist; SEM, standard error of the mean.
<sup>a</sup> Significantly differed between subgroups.

### *fc*MRI Acquisition and Processing

Magnetic resonance imaging (MRI) data were acquired on a 3T magnet. Each session included 5 high-resolution structural T1 (voxel size = 0.8 × 0.8 × 0.8 mm; TR = 2200 ms; TE = 2.83 ms), 6 functional MRI (rest, script imagery, Stroop, Hariri, Nback, CO2; voxel size = 2 × 2 × 3 mm; TR = 3000 ms; TE = 30 ms), 1 diffusion weighted imaging (DWI), 1 arterial spin labeling (ASL), and 1 FLAIR scan (voxel size = 0.8 × 0.8 × 0.8 mm; TR = 2200 ms; TE = 2.83 ms). Here, we processed and investigated functional connectivity at rest (10 minutes; 200 frames) and during symptom provocation (12 minutes; 240 frames). T1 and FLAIR scans were used for tissue segmentation and coregistration.

The symptom-provocation task consisted of subject-specific neutral and trauma scripts (1 minute each) based on a structured questionnaire completed during the interview with each participant. The script was recorded in second person (i.e., “You are in Iraq…”) and in a voice sex-matched to the participant. During scanning, the script was played for 60 seconds, and participants were instructed to “recall and relive the experience.” For an additional 60 seconds, the participants were instructed to “think about the events” and recreate the experience. Then, they were instructed to “let it go” for an additional 60 seconds. These 3-stage retelling and clearing instructions were repeated 4 times, alternating between neutral and trauma scripts. Considering the potential for carry over and to obtain stable functional connectivity estimates, the full 12-minute run was used for computing GBCr during symptom provocation. The Human Connectome Pipeline was adapted to conduct surface-based preprocessing and optimize registration (31). Details of our image processing pipeline were previously reported (26) and are provided in the Supplemental Information. GBCr calculation followed our previous reports (17,18,26,28); i.e., they were computed as the average of the correlations between each voxel and all other voxels in the brain gray matter (see Supplemental Information).

### Statistical Analyses

Statistical Package for the Social Sciences (SPSS, version 24) was used for the behavioral and region of interest (ROI) analyses. The distribution of outcome measures was examined using probability plots and test statistics. Transformations and nonparametric tests were used as necessary. Estimates of variation are provided as the standard error of the mean (SEM). Significance was set at *P* ≤ .05, with 2-tailed tests. ANOVA and chi-squares were used to compare behavioral data across groups. Body mass index (BMI), intelligence quotient (IQ), and ethnicity differed between the study groups. Therefore, these variables were included as covariates in the vertex/voxel-wise and ROI analyses.

Vertex/voxel-wise *fc*MRI nonparametric analyses used FSL Permutation Analysis of Linear Models (PALM), with tail approximation and cluster mass threshold of 1.96 for Type I error correction (corrected *α* = .05) (32). First, we conducted a data-driven, whole-brain analysis using independent *t* tests to identify clusters with altered GBCr in the PTSD group compared to all controls, at rest and during symptom provocation. Then, we extracted the identified clusters (vertex/voxel *P* < .005; corrected *α* = .05) and conducted follow-up ROI analyses to better characterize the GBCr abnormalities across tasks and subgroups. This was accomplished by constructing a general linear model (GLM) that examined the effects of groups (PTSD vs. CC vs. HC), tasks (rest vs. scripts), and group*tasks interaction, followed by *post hoc* pairwise comparisons. Finally, we conducted exploratory linear regression analysis examining whether at-rest salience GBCr predicts GBCr during symptom provocation.

## RESULTS

Participants were well matched for age, sex, and race (Table 1). Scans were excluded if they had any frame with absolute motion > 2 mm, any frame with relative motion > 2 mm, or more than 50% of frames with frame displacement > 0.3 mm. Accordingly, 5 rest and 6 script scans were excluded.

### Whole-Brain Data-Driven Analyses: Disrupted Connectivity Within the Salience Networks

During symptom provocation, the whole-brain analysis revealed a significantly higher GBCr in PTSD participants compared to controls in areas within the right hemisphere, including the anterior insula, caudal- ventrolateral prefrontal, and rostral-ventrolateral parietal cortices (Fig. 1A; orange-yellow clusters). Notably, these areas primarily overlap with the ventral and dorsal salience networks (Fig. 1B; orange and blue clusters) (6). Moreover, we found significant clusters of lower GBCr in the caudal-dorsal areas of the cerebellum in participants with PTSD compared to control (Fig. 2; blue clusters). At rest, we found no significant differences in GBCr between the study groups.

**Figure 1.**
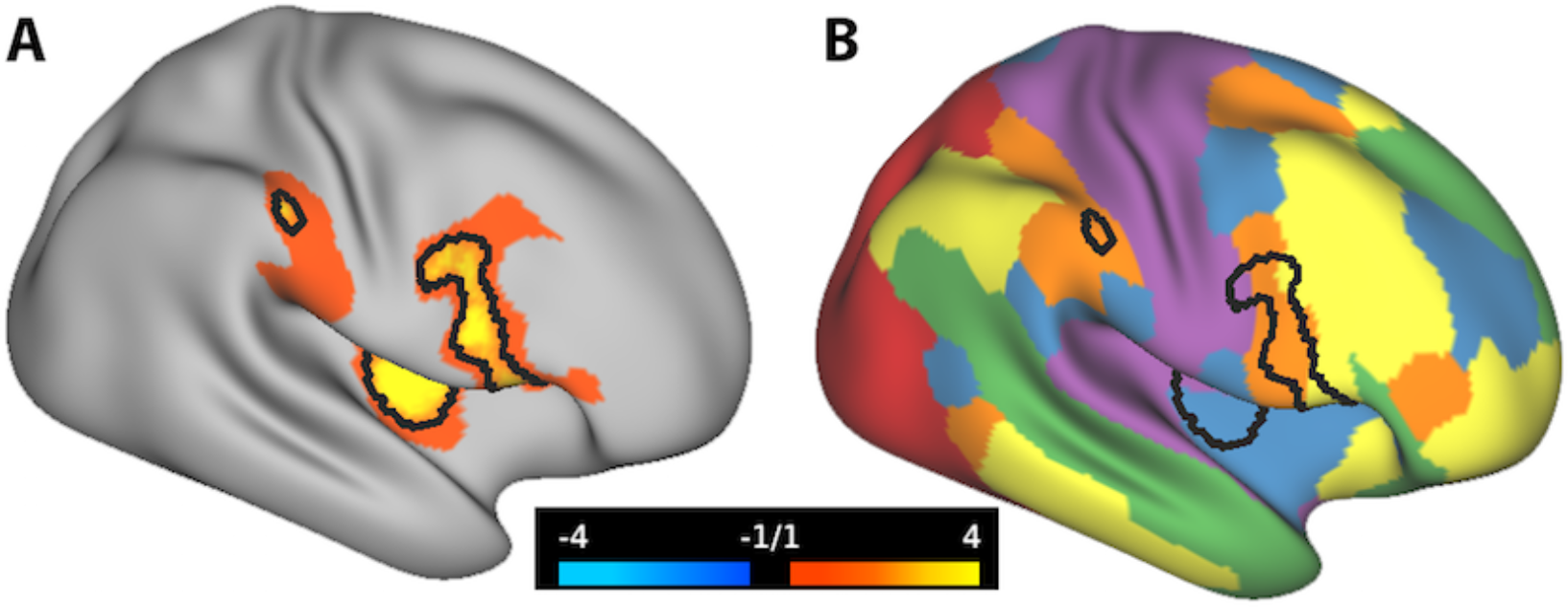
Cortical Global Connectivity in US Army Soldiers with Posttraumatic Stress Disorder (PTSD) **A.** The orange-yellow clusters mark the vertices with increased global brain connectivity with global signal regression (GBCr) in PTSD compared to controls during symptom provocation. The black lines mark the vertices with *P* < .005 and corrected *α* = .05. **B.**The map of 6 intrinsic connectivity networks: ventral salience (blue), dorsal salience (orange), central executive (yellow), default mode (green), visual (red), and sensorimotor (purple). The black lines in B mirror the black lines in A.

**Figure 2.**
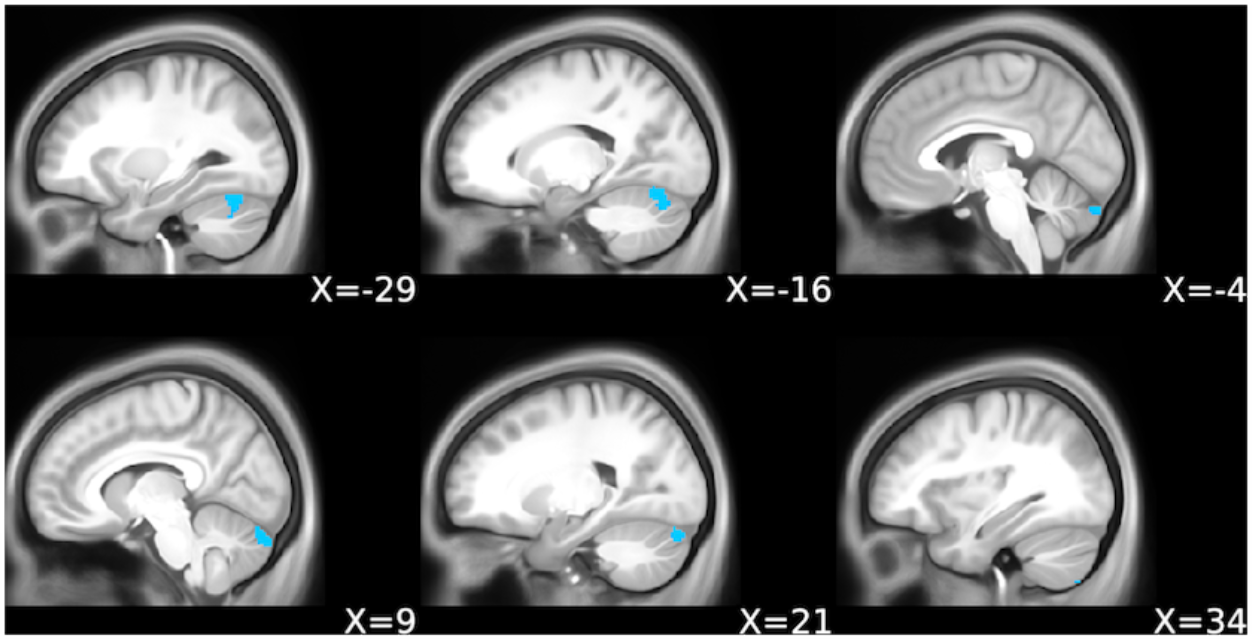
Cerebellar Global Connectivity in US Army Soldiers with Posttraumatic Stress Disorder (PTSD) The blue clusters mark the voxels with reduced global brain connectivity with global signal regression (GBCr) in PTSD compared to controls during symptom provocation (*P* < .005 and corrected *α* = .05).

### ROI Post hoc Analyses: Increased Connectivity During Symptom Provocation in PTSD

At rest and during symptom provocation, we extracted average GBCr from each subject within 2 ROIs. The salience ROI included areas that showed significantly high GBCr in PTSD (Fig. 1). The cerebellar ROI included areas with low GBCr in PTSD (Fig. 2). Investigating the salience ROI, the GLM showed a significant group effect (*F*_(2,85)_ = 7.6, *P* = .001; Fig. 3), with higher salience GBCr in PTSD compared to CC (*P* =.001) and HC (*P* = .003), but no differences between HC and CC (*P* = .71). We also found a significant group*task interaction (*F*_(2,85)_ = 4.4, *P* = .01; Fig. 4), with significant increase in salience GBCr during symptom provocation compared to rest in PTSD (*P* = .04), but not in CC (*P* = .13) and HC (*P* = .09).

**Figure 3.**
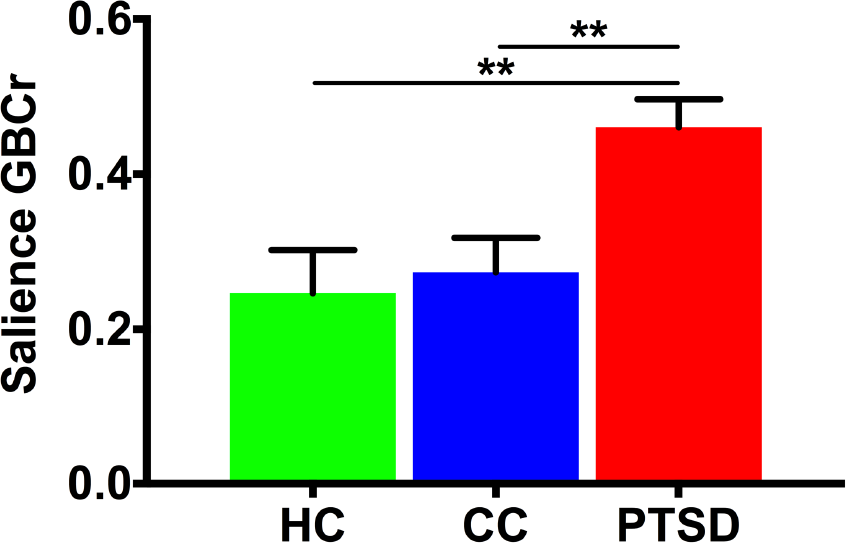
Overall Salience Global Connectivity in US Army Soldiers with Posttraumatic Stress Disorder (PTSD) There was a significant main group effect with increased overall (i.e., at rest and during trauma recollection) global brain connectivity with global signal regression (GBCr) in PTSD compared to combat (CC) and healthy controls (HC). ** *P* ≤ .01.

**Figure 4.**
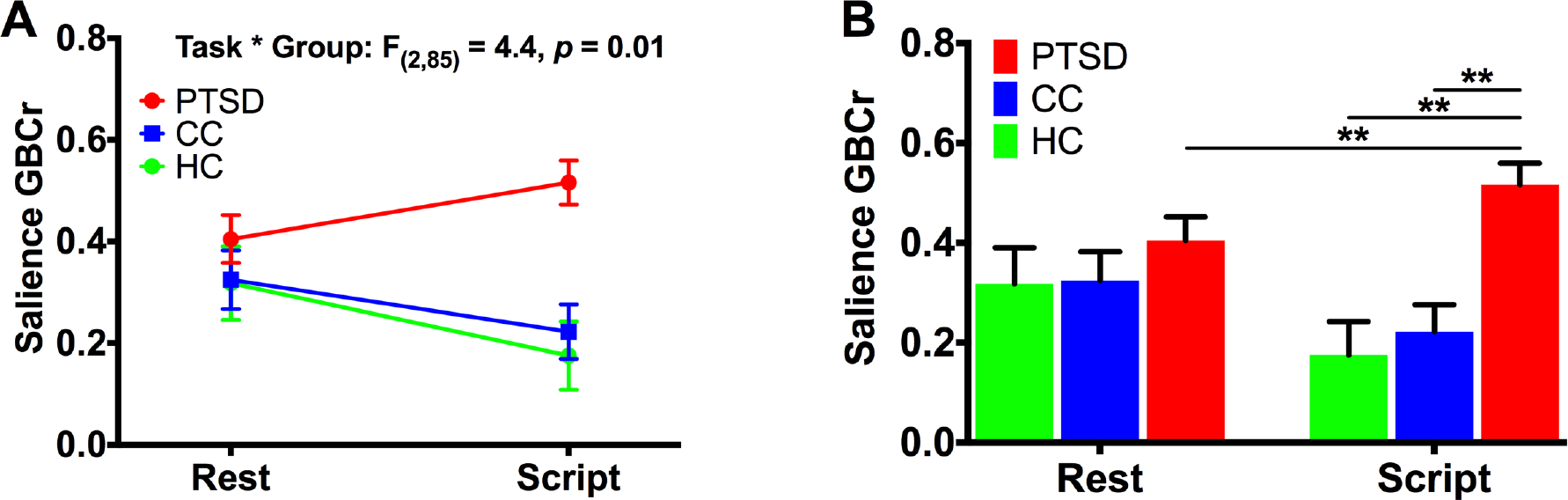
Effect of Symptom Provocation on Salience Global Connectivity in US Army Soldiers with Posttraumatic Stress Disorder (PTSD) A. There was a significant group by task interaction effect on salience global brain connectivity with global signal regression (GBCr). **B.** *Post hoc* comparison shows significant increase in GBCr during trauma recollection (i.e., script imagery) compared to during resting state in PTSD but not in controls. The higher GBCr values in PTSD compared to combat (CC) and healthy controls (HC) were significant only during trauma recollection, but not at rest. ** *P* ≤ .01. Rest indicates resting state; Script, script imagery.

Although salience GBCr differed between groups only during symptom provocation, it was numerically higher in PTSD even at rest (Fig. 4B), raising the question as to whether a subgroup of PTSD subjects experienced symptoms and engaged the salience network at rest. Therefore, we conducted a linear regression examining whether salience GBCr at rest predicts GBCr during symptom provocation. We found that salience GBCr at rest positively predicted salience GBCr during symptom provocation in the PTSD [*F*_(1,42)_ = 16.0, *P* < .001], but not in the CC [*F*_(1,25)_ = 1.1, *P* = .49] and HC groups [*F*_(1,19)_ = 0.5, *P* = .31].

In the cerebellar ROI, we found a significant group effect (*P* = 0.02), with lower cerebellar GBCr in PTSD compared to CC (*P* = .006) but not HC (*P* = .08) (Fig. S1). There were no significant effects of task (*P* = .08), or group*task interaction (*P* = .08), and no difference between CC and HC (*P* = .61).

## DISCUSSION

The data-driven results identified widespread disruption of functional connectivity in areas within the salience network of US Army soldiers suffering from PTSD. The salience dysconnectivity was evident during symptom provocation but not at rest. The increase in salience global connectivity was found in the PTSD group compared to both combat and healthy controls. Compared to connectivity at rest, symptom provocation induced significant increase in salience global connectivity in the PTSD group but not the control groups. In addition, the data-driven analysis revealed reduced global connectivity in PTSD in areas within the posterior lobe of the cerebellum, a cerebellar region that primarily serves cognitive functions. The reduction in cerebellar connectivity was significant in PTSD compared to combat but not to healthy controls. The identified clusters of salience and cerebellar connectivity did not differ between combat and healthy controls. Finally, although the salience connectivity did not differ between groups at rest, PTSD patients have shown numerically higher values that significantly predicted increased global connectivity during symptom provocation. The latter findings raise the possibility that the identified salience dysconnectivity may reflect a state-dependent abnormality that is concealed at rest but more evident in the presence of trauma reminders.

The present investigation extends past research by employing a data-driven approach rather than a seed-based analysis and by examining global connectivity at rest and during symptom provocation. Previous studies have reported greater functional connectivity between salience network regions in individuals with PTSD compared to controls (33–36). The salience network is believed to play a critical role in (a) threat detection, (b) arbitration between other large-scale intrinsic networks, particularly the default mode and central executive networks, and (c) modulation of autonomic reactivity to salient stimuli (37). PTSD is associated with impairment in each of these 3 salience-related functions, i.e., increased threat detection, impaired default and executive connectivity, and autonomic dysregulation (1,4,38). A recent network-based framework posits that heightened salience connectivity in PTSD may result in impaired intrinsic connectivity related to the arbitration function, leading to low threshold for attention to trauma-related cues, which may help explain hyperarousal symptoms in PTSD patients (3).

Overall, the study findings support our *a priori* hypothesis of state-dependent symptom-induced increases in cortical GBCr, putatively concealing any GBCr deficits related to synaptic loss. Reduced prefrontal GBCr has been directly associated with trauma- and stress-related synaptic loss (1,27). Moreover, these cortical GBCr deficits have been reported in several psychiatric disorders with a considerable chronic stress component, including unipolar and bipolar depression, obsessive-compulsive disorder, and schizophrenia (17–25). Intriguingly, despite extensive evidence suggesting stress- and PTSD-related cortical synaptic loss (39,40), the current study and previous report (28) failed to demonstrate reduction in cortical GBCr in PTSD. Considering previous evidence correlating reduced cortical GBCr with avoidance and numbing symptoms, and increased cortical GBCr with arousal symptoms (28), we interpret the increase of GBCr during trauma recollection as evidence that the functional dysconnectivity related to synaptic loss may have been masked by superimposed increased connectivity related to arousal and perhaps reexperiencing symptoms.

Unexpectedly, the identified cortical GBCr abnormalities showed a remarkable spatial pattern with noticeable overlap with the dorsal and ventral salience networks, interestingly limited to the right hemisphere. Lateralization in the anterior insula, an essential node within the salience network, has long been suggested with the right insula highly connected to the sympathetic autonomic system and plays a main role in attentional orientation to salient stimuli, interoception, and arousal (41). In contrast, the left hemisphere insula is more connected to the parasympathetic system and is associated with functions such as nourishment, gesture, positive affect, and cognitive and affective control (41). Importantly, PTSD is associated with an overactive sympathetic nervous system and attentional orientation to salient stimuli including heightened threat detection (38,42,43). The localization of the study findings to the salience network and right hemisphere during trauma recollection further underscores the neurobiological and clinical relevance of functional nodal strength as measured by GBCr. It also highlights the utility of longitudinal connectivity designs, where connectivity is tested at rest as well as during or following an intervention.

Another unpredicted finding is the GBCr reduction in the posterior lobe of the cerebellum during symptom provocation. While the cerebellum has long been viewed as a unit of motor control, it is becoming increasingly apparent that the cerebellum, especially the neocerebellum (posterior lobe) that is mainly connected to the cerebral cortex, plays important roles in emotion and cognition (44). Recent work by Holmes and colleagues also found reduced nodal strength in the cerebellum of PTSD patients, as well as reduced volume and structural covariance (45). Other studies have also reported structural and functional abnormalities in the cerebellum in PTSD (46–51). Importantly, the cerebellar abnormalities in the current study were evident only during symptom provocation. This finding supports the presence of a direct relationship between reduced cerebellar global connectivity and trauma recollection. In addition, it shows the potential utility of paradigm-based connectivity to induce homogenous and exaggerated effects, which facilitate the interpretation of the results and may enhance the reproducibility of the findings in future studies.

Finally, in contrast to a previous report of reduced GBCr in the anterior hippocampus in veterans with PTSD (28), the current study failed to show any subcortical dysconnectivity at rest or during symptom provocation. A putative explanation is that PTSD may be associated with a dual pathology, including a subgroup of patients suffering from amino-acid pathology (ABP) and another subgroup having primarily a monoamine pathology (MBP) (1). The ABP characteristics include reduced gray matter (e.g., hippocampal volume reduction), reduced amino-acid levels (e.g., low glutamate), increased inflammation, hypothalamic-pituitary axis dysregulation, and chronic, complex and treatment-resistant PTSD (1). Therefore, we speculate that the difference in study population may have led to the differing results. In particular, the target population and study criteria of the previous report may have led to enrolling patients with ABP, as evidenced by widespread structural and connectivity deficits in the previous cohort (28,52–55).

### Limitations and Strengths

The experimental within-session design directly associated trauma recollection with increased salience connectivity in individuals suffering from PTSD. However, we cannot determine whether the salience disturbance predates the development of PTSD or is a consequence of PTSD pathology. Predeployment longitudinal studies would be critical to ascertain the causal relationship. An alternative approach, to shed light on the relationship between salience network and PTSD, is to determine whether these salience disturbances will normalize following successful treatment, an investigation that will be reported elsewhere. Another limitation is that the current study is mainly investigating the PTSD group. Therefore, to optimize type I and type II errors, the study did not fully assess for differences between combat and healthy controls. In that regard, it is important to underscore that the *post hoc* ROI analyses conducted are dependent on the whole-brain investigation and should be viewed and interpreted only within the context of better characterizing the data-driven results. Finally, the timing (i.e., symptom-induced), localization, and lateralization of the disturbance in nodal strength strongly implicate the ventral and dorsal salience networks. However, the nodal strength measured is based on whole-brain global connectivity and not limited to a specific ICN. Therefore, the identified salience dysconnectivity may reflect increased internal (i.e., within network) and/or external connectivity (i.e., between networks). Future studies may employ network-restricted topology approaches to further investigate the role of ICNs in the pathology of PTSD (53).

The strengths of the study include (a) a relatively large sample in a less-often investigated target population (i.e., active duty military personnel); (b) the use of state-of-the-art neuroimaging methods based on the Human Connectome Pipeline, including enhanced registration, surface-based analysis, and nonparametric correction for vertex/voxel-wise multiple comparisons; (c) the use of GBCr, a well-validated measure of nodal strength that has been repeatedly associated with psychopathology and successful treatment, and that does not require *a priori* selection of seed or ROI; and (d) the use of trauma recollection design to identify symptom-specific network disturbances.

## CONCLUSIONS

The results strongly implicate the salience network in the pathophysiology of PTSD, demonstrating symptom-induced lateralized increase of global connectivity in brain areas within the ventral and dorsal salience networks in the right hemisphere. The high salience global connectivity during trauma recollection was evident in PTSD compared to both combat and healthy controls. Although there was no significant dysconnectivity at rest, the salience global connectivity during trauma recollection was positively predicted by salience connectivity at rest. Together, the results support the hypothesis that the reduction of cortical global connectivity due to synaptic loss in PTSD may be concealed by state-dependent, arousal-related increases in connectivity. Finally, the study found reduced global connectivity in the posterior lobe of the cerebellum, contributing to accumulating evidence implicating this essential brain region in the pathology of PTSD.

## ACKNOWLEDGMENTS

The authors would like to thank the individuals who participated in these studies for their invaluable contribution. Funding for this work was made possible by grants to the STRONG STAR Consortium by the U.S. Department of Defense through the U.S. Army Medical Research and Materiel Command, Congressionally Directed Medical Research Programs, Psychological Health and Traumatic Brain Injury Research Program awards W81XWH-08-02-109 (Alan Peterson), W81XWH-08-02-0112 (Peter Fox), W81XWH-08-02-0114 (Brett Litz), and W81XWH-08-02-0116 (Patricia Resick). Some of the investigators also had additional support from the National Institute of Mental Health (K23MH101498) and the VA National Center for PTSD. The views expressed in this article are solely those of the authors and do not represent and endorsement by or the official policy or position of the Department of Defense, the Department of Veterans Affairs, the National Institutes of Health, or the US Government.

## SUPPLEMENTAL INFORMATION

### Image Processing

The Human Connectome Project (HCP) Pipelines (github.com/Washington-University/Pipelines) were adapted to process the imaging data (1). Briefly, the adapted minimal preprocessing included FreeSurfer automatic segmentation and parcellation of high- resolution structural scans, deletion of the first 5 volumes, slice timing correction, motion correction, intensity normalization, brain masking, and registration of *f*MRI images to structural MRI and standard template, while minimizing smoothing from interpolation. Then, the cortical gray matter ribbon voxels and each subcortical parcel were projected to a standard Connectivity Informatics Technology Initiative (CIFTI) 2mm grayordinate space. ICA-FIX was run to identify and remove artifacts (2,3), followed by mean grayordinate time series regression (MGTR; which is comparable to global signal regression in volume data). The latter two processing steps (FIX+MGTR) have been found to significantly reduce motion-correlated artifacts (4). In addition, there were no differences (*P* > .1) in head motion during *f*MRI session between the study groups at rest (mean ±*SEM*; PTSD = 0.09 ±0.009; CC = 0.07 ±0.003; HC = 0.09 ±0.013) and during symptoms provocation (mean ±*SEM*; PTSD = 0.10 ±0.008; CC = 0.09 ±0.009; HC = 0.10 ±0.012).

Details of global brain connectivity with global signal regression (GBCr) methods were previously described (5–16). Briefly, time series were demeaned and normalized, followed by generating dense connectomes correlating each vertex/voxel with all other vertices/voxels in the CIFTI grayordinates, and then transformed to Fisher *z* values. For each vertex/voxel, GBCr is calculated as the standardized (*z* scored) average across those Fisher *z* values with parcel-constrained smoothing (sigma = 4.2 mm), which generates a map for each *f*MRI session where each vertex/voxel value represents the functional connectivity strength of that grayordinate with the rest of the brain. In graph theory terms, GBCr (also known as functional connectivity strength; FCS [17)]) is considered a weighted measure of nodal strength of a voxel in the whole brain network — determining brain hubs and examining the coherence between a local region and the rest of the brain (18).

**Figure S1.**
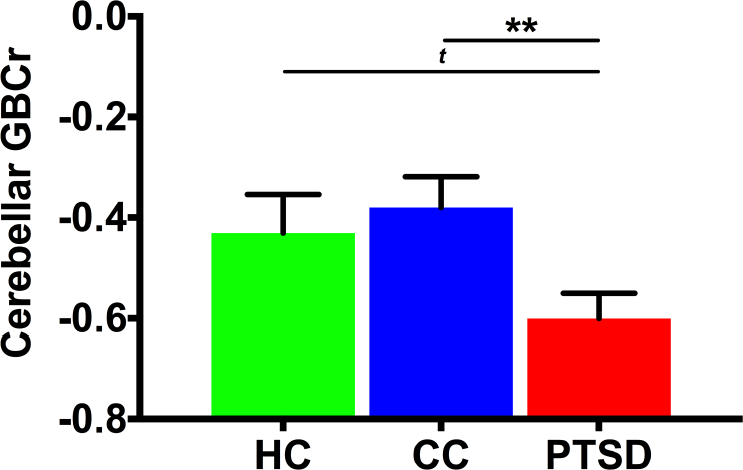
Overall Cerebellar Global Connectivity in US Army Soldiers with Posttraumatic Stress Disorder (PTSD). There was a significant main group effect with increased overall (i.e., at rest and during trauma recollection) global brain connectivity with global signal regression (GBCr) in PTSD compared to combat controls (CC), with a trend level significance, compared to healthy control (HC). ** *P* ≤ .01; ^*t*^ *P* ≤ .1.

Similar to previous studies (5–11,13,17), we have used GBCr, instead of GBC without global signal regression (GBCnr), because the study hypotheses were based on previous GBCr findings (5–7), which provided the rationale for the current report and will facilitate the interpretation of the study findings. In addition, previous work underscored the need for MGTR to adequately minimize spurious artifacts (4).

## REFERENCES

1. Abdallah CG, Averill LA, Akiki TJ, Raza M, Averill CL, Gomaa H, et al. (2019): The neurobiology and pharmacotherapy of posttraumatic stress disorder. Annu Rev Pharmacol Toxicol 59:17.1–17.19. doi:10.1146/annurev-pharmtox-010818-021701

2. Krystal JH, Davis LL, Neylan TC, Raskind MA, Schnurr PP, Stein MB, et al. (2017): It is time to address the crisis in the pharmacotherapy of posttraumatic stress disorder: A consensus statement of the PTSD Psychopharmacology Working Group. Biol Psychiatry 82:e51–e59. doi:10.1016/j.biopsych.2017.03.007

3. Akiki TJ, Averill CL, Abdallah CG (2017): A network-based neurobiological model of PTSD: Evidence from structural and functional neuroimaging studies. Curr Psychiatry Rep 19:81. doi:10.1007/s11920-017-0840-4

4. Sheynin J, Liberzon I (2017): Circuit dysregulation and circuit-based treatments in posttraumatic stress disorder. Neurosci Lett 649:133–138.

5. Biswal B, Yetkin FZ, Haughton VM, Hyde JS (1995): Functional connectivity in the motor cortex of resting human brain using echo-planar MRI. Magn Reson Med 34:537–541.

6. Akiki TJ, Abdallah CG (2018): Determining the hierarchical architecture of the human brain using subject-level clustering of functional networks [Manuscript on preprint server]. bioRxiv. doi:10.1101/350462

7. Power JD, Cohen AL, Nelson SM, Wig GS, Barnes KA, Church JA, et al. (2011): Functional network organization of the human brain. Neuron 72:665–678.

8. Yeo BT, Krienen FM, Sepulcre J, Sabuncu MR, Lashkari D, Hollinshead M, et al. (2011): The organization of the human cerebral cortex estimated by intrinsic functional connectivity. J Neurophysiol 106:1125–1165.

9. Menon V (2011): Large-scale brain networks and psychopathology: A unifying triple network model. Trends Cogn Sci 15:483–506.

10. Cole MW, Bassett DS, Power JD, Braver TS, Petersen SE (2014): Intrinsic and task-evoked network architectures of the human brain. Neuron 83:238–251.

11. Larson-Prior LJ, Zempel JM, Nolan TS, Prior FW, Snyder AZ, Raichle ME (2009): Cortical network functional connectivity in the descent to sleep. Proc Natl Acad Sci U S A 106:4489–4494.

12. Bullmore E, Sporns O (2009): Complex brain networks: Graph theoretical analysis of structural and functional systems. Nat Rev Neurosci 10:186–198.

13. Cole MW, Pathak S, Schneider W (2010): Identifying the brain’s most globally connected regions. NeuroImage 49:3132–3148.

14. Cole MW, Yarkoni T, Repovs G, Anticevic A, Braver TS (2012): Global connectivity of prefrontal cortex predicts cognitive control and intelligence. J Neurosci 32:8988–8999.

15. Liang X, Connelly A, Calamante F (2014): Graph analysis of resting-state ASL perfusion MRI data: nonlinear correlations among CBF and network metrics. Neuroimage 87:265–275.

16. Liang X, Zou Q, He Y, Yang Y (2013): Coupling of functional connectivity and regional cerebral blood flow reveals a physiological basis for network hubs of the human brain. Proc Natl Acad Sci U S A 110:1929–1934.

17. Abdallah CG, Averill CL, Salas R, Averill LA, Baldwin PR, Krystal JH, et al. (2017): Prefrontal connectivity and glutamate transmission: relevance to depression pathophysiology and ketamine treatment. Biol Psychiatry Cogn Neurosci Neuroimaging 2:566–574.

18. Abdallah CG, Averill LA, Collins KA, Geha P, Schwartz J, Averill C, et al. (2017): Ketamine treatment and global brain connectivity in major depression. Neuropsychopharmacology 42:1210–1219.

19. Murrough JW, Abdallah CG, Anticevic A, Collins KA, Geha P, Averill LA, et al. (2016): Reduced global functional connectivity of the medial prefrontal cortex in major depressive disorder. Hum Brain Mapp 37:3214–3223.

20. Scheinost D, Holmes SE, DellaGioia N, Schleifer C, Matuskey D, Abdallah CG, et al. (2018): Multimodal investigation of network level effects using intrinsic functional connectivity, anatomical covariance, and structure-to-function correlations in unmedicated major depressive disorder. Neuropsychopharmacology 43:1119–1127.

21. Wang L, Dai Z, Peng H, Tan L, Ding Y, He Z, et al. (2014): Overlapping and segregated resting-state functional connectivity in patients with major depressive disorder with and without childhood neglect. Hum Brain Mapp 35:1154–1166.

22. Anticevic A, Brumbaugh MS, Winkler AM, Lombardo LE, Barrett J, Corlett PR, et al. (2013): Global prefrontal and fronto-amygdala dysconnectivity in bipolar I disorder with psychosis history. Biol Psychiatry 73:565–573.

23. Anticevic A, Hu S, Zhang S, Savic A, Billingslea E, Wasylink S, et al. (2014): Global resting-state functional magnetic resonance imaging analysis identifies frontal cortex, striatal, and cerebellar dysconnectivity in obsessive-compulsive disorder. Biol Psychiatry 75:595–605.

24. Anticevic A, Corlett PR, Cole MW, Savic A, Gancsos M, Tang Y, et al. (2015): N-methyl-D-aspartate receptor antagonist effects on prefrontal cortical connectivity better model early than chronic schizophrenia. Biol Psychiatry 77:569–580.

25. Cole MW, Anticevic A, Repovs G, Barch D (2011): Variable global dysconnectivity and individual differences in schizophrenia. Biol Psychiatry 70:43–50.

26. Abdallah CG, Dutta A, Averill CL, McKie S, Akiki TJ, Averill LA, et al. (2018): Ketamine, but not the NMDAR antagonist lanicemine, increases prefrontal global connectivity in depressed patients. Chronic Stress 2:2470547018796102. doi:10.1177/2470547018796102

27. Abdallah CG, Sanacora G, Duman RS, Krystal JH (2018): The neurobiology of depression, ketamine and rapid-acting antidepressants: Is it glutamate inhibition or activation? Pharmacol Ther 190:148–158.

28. Abdallah CG, Wrocklage KM, Averill CL, Akiki T, Schweinsburg B, Roy A, et al. (2017): Anterior hippocampal dysconnectivity in posttraumatic stress disorder: A dimensional and multimodal approach. Transl Psychiatry 7:e1045. doi:10.1038/tp.2017.12

29. Driesen NR, McCarthy G, Bhagwagar Z, Bloch MH, Calhoun VD, D’Souza DC, et al. (2013): The impact of NMDA receptor blockade on human working memory-related prefrontal function and connectivity. Neuropsychopharmacology 38:2613–2622.

30. Driesen NR, McCarthy G, Bhagwagar Z, Bloch M, Calhoun V, D’Souza DC, et al. (2013): Relationship of resting brain hyperconnectivity and schizophrenia-like symptoms produced by the NMDA receptor antagonist ketamine in humans. Mol Psychiatry 18:1199–1204.

31. Glasser MF, Sotiropoulos SN, Wilson JA, Coalson TS, Fischl B, Andersson JL, et al. (2013): The minimal preprocessing pipelines for the Human Connectome Project. Neuroimage 80:105–124.

32. Winkler AM, Ridgway GR, Webster MA, Smith SM, Nichols TE (2014): Permutation inference for the general linear model. Neuroimage 92:381–397.

33. Rabinak CA, Angstadt M, Welsh RC, Kenndy AE, Lyubkin M, Martis B, et al. (2011): Altered amygdala resting-state functional connectivity in post-traumatic stress disorder. Front Psychiatry 2:62. doi:10.3389/fpsyt.2011.00062

34. Sripada RK, King AP, Garfinkel SN, Wang X, Sripada CS, Welsh RC, et al. (2012): Altered resting-state amygdala functional connectivity in men with posttraumatic stress disorder. J Psychiatry Neurosci 37:241–249.

35. Sripada RK, King AP, Welsh RC, Garfinkel SN, Wang X, Sripada CS, et al. (2012): Neural dysregulation in posttraumatic stress disorder: evidence for disrupted equilibrium between salience and default mode brain networks. Psychosom Med 74:904–911.

36. Thome J, Frewen P, Daniels JK, Densmore M, Lanius RA (2014): Altered connectivity within the salience network during direct eye gaze in PTSD. Borderline Personal Disord Emot Dysregul. 1:17. doi:10.1186/2051-6673-1-17

37. Menon V, Uddin LQ (2010): Saliency, switching, attention and control: A network model of insula function. Brain Struct Funct 214:655–667.

38. Pitman RK, Rasmusson AM, Koenen KC, Shin LM, Orr SP, Gilbertson MW, et al. (2012): Biological studies of post-traumatic stress disorder. Nat Rev Neurosci 13:769–787.

39. Averill LA, Abdallah CG, Pietrzak RH, Averill CL, Southwick SM, Krystal JH, Harpaz-Rotem I (2017): Combat exposure severity is associated with reduced cortical thickness in combat veterans: A preliminary report. Chronic Stress 1:2470547017724714. doi:10.1177/2470547017724714

40. Wrocklage KM, Averill LA, Cobb Scott J, Averill CL, Schweinsburg B, Trejo M, et al. (2017): Cortical thickness reduction in combat exposed US veterans with and without PTSD. Eur Neuropsychopharmacol 27:515–525.

41. Montalbano MJ, Tubbs RS (2018): Lateralization of the insular cortex. In: Turgut M, Yurttas C, Tubbs RS, editors. Island of Reil (Insula) in the Human Brain. New York: Springer, pp 129–132.

42. Yehuda R (2002): Post-traumatic stress disorder. N Engl J Med 346:108–114.

43. Liberzon I, Abelson JL (2016): Context processing and the neurobiology of post-traumatic stress disorder. Neuron 92:14–30.

44. Schutter DJ, van Honk J (2005): The cerebellum on the rise in human emotion. Cerebellum 4:290–294.

45. Holmes SE, Scheinost D, DellaGioia N, Davis MT, Matuskey D, Pietrzak RH, et al. (2018): Cerebellar and prefrontal cortical alterations in PTSD: structural and functional evidence. Chronic Stress 2:2470547018786390. doi:10.1177/2470547018786390

46. Hayes JP, Hayes SM, Mikedis AM (2012): Quantitative meta-analysis of neural activity in posttraumatic stress disorder. Biol Mood Anxiety Disord 2:9. doi:10.1186/2045-5380-2-9

47. Koch SB, van Zuiden M, Nawijn L, Frijling JL, Veltman DJ, Olff M (2016): Aberrant resting-state brain activity in posttraumatic stress disorder: A meta-analysis and systematic review. Depress Anxiety 33:592–605.

48. Baldacara L, Borgio JGF, Araujo C, Nery-Fernandes F, Lacerda ALT, Moraes W, et al. (2012): Relationship between structural abnormalities in the cerebellum and dementia, posttraumatic stress disorder and bipolar disorder. Dement Neuropsychol 6:203–211.

49. Baldacara L, Jackowski AP, Schoedl A, Pupo M, Andreoli SB, Mello MF, et al. (2011): Reduced cerebellar left hemisphere and vermal volume in adults with PTSD from a community sample. J Psychiatr Res 45:1627–1633.

50. De Bellis MD, Kuchibhatla M (2006): Cerebellar volumes in pediatric maltreatment-related posttraumatic stress disorder. Biol Psychiatry 60:697-703.

51. Rabellino D, Densmore M, Theberge J, McKinnon MC, Lanius RA (2018): The cerebellum after trauma: resting-state functional connectivity of the cerebellum in posttraumatic stress disorder and its dissociative subtype. Hum Brain Mapp 39:3354–3374.

52. Akiki TJ, Averill CL, Wrocklage KM, Schweinsburg B, Scott JC, Martini B, et al. (2017): The association of PTSD symptom severity with localized hippocampus and amygdala abnormalities. Chronic Stress 1:2470547017724069. doi:10.1177/2470547017724069

53. Akiki TJ, Averill CL, Wrocklage KM, Scott JC, Averill LA, Schweinsburg B, et al. (2018): Default mode network abnormalities in posttraumatic stress disorder: A novel network-restricted topology approach. Neuroimage. 176:489–498.

54. Averill CL, Averill LA, Wrocklage KM, Scott JC, Akiki TJ, Schweinsburg B, et al. (2018): Altered white matter diffusivity of the cingulum angular bundle in posttraumatic stress disorder. Mol Neuropsychiatry 4:75–82.

55. Averill CL, Satodiya RM, Scott JC, Wrocklage KM, Schweinsburg B, Averill LA, et al. (2017): Posttraumatic stress disorder and depression symptom severities are differentially associated with hippocampal subfield volume loss in combat veterans. Chronic Stress 1:2470547017744538. doi:10.1177/2470547017744538

## REFERENCES

1. Glasser MF, Sotiropoulos SN, Wilson JA, Coalson TS, Fischl B, Andersson JL, et al. (2013): The minimal preprocessing pipelines for the Human Connectome Project. Neuroimage. 80:105–124.

2. Salimi-Khorshidi G, Douaud G, Beckmann CF, Glasser MF, Griffanti L, Smith SM (2014): Automatic denoising of functional MRI data: Combining independent component analysis and hierarchical fusion of classifiers. Neuroimage. 90:449–468.

3. Griffanti L, Salimi-Khorshidi G, Beckmann CF, Auerbach EJ, Douaud G, Sexton CE, et al. (2014): ICA-based artefact removal and accelerated fMRI acquisition for improved resting state network imaging. Neuroimage. 95:232–247.

4. Burgess GC, Kandala S, Nolan D, Laumann TO, Power JD, Adeyemo B, et al. (2016): Evaluation of denoising strategies to address motion-correlated artifacts in resting-state functional magneticresonance imaging data from the Human Connectome Project. Brain Connect 6:669–680.

5. Abdallah CG, Averill CL, Salas R, Averill LA, Baldwin PR, Krystal JH, et al. (2017): Prefrontal connectivity and glutamate transmission: Relevance to depression pathophysiology and ketamine treatment. Biol Psychiatry Cogn Neurosci Neuroimaging 2:566–574.

6. Abdallah CG, Averill LA, Collins KA, Geha P, Schwartz J, Averill C, et al. (2017): Ketamine treatment and global brain connectivity in major depression. Neuropsychopharmacology. 42:1210–1219.

7. Abdallah CG, Wrocklage KM, Averill CL, Akiki T, Schweinsburg B, Roy A, et al. (2017): Anterior hippocampal dysconnectivity in posttraumatic stress disorder: A dimensional and multimodal approach. Transl Psychiatry 7:e1045. doi:10.1038/tp.2017.12

8. Murrough JW, Abdallah CG, Anticevic A, Collins KA, Geha P, Averill LA, et al. (2016): Reduced global functional connectivity of the medial prefrontal cortex in major depressive disorder. Hum Brain Mapp 37:3214–3223.

9. Anticevic A, Brumbaugh MS, Winkler AM, Lombardo LE, Barrett J, Corlett PR, et al. (2013): Global prefrontal and fronto-amygdala dysconnectivity in bipolar I disorder with psychosis history. Biol Psychiatry. 73:565–573.

10. Anticevic A, Corlett PR, Cole MW, Savic A, Gancsos M, Tang Y, et al. (2015): N-methyl-D-aspartate receptor antagonist effects on prefrontal cortical connectivity better model early than chronic schizophrenia. Biol Psychiatry 77:569–580.

11. Anticevic A, Hu S, Zhang S, Savic A, Billingslea E, Wasylink S, et al. (2014): Global resting-state functional magnetic resonance imaging analysis identifies frontal cortex, striatal, and cerebellar dysconnectivity in obsessive-compulsive disorder. Biol Psychiatry 75:595–605.

12. Anticevic A, Hu X, Xiao Y, Hu J, Li F, Bi F, et al. (2015): Early-course unmedicated schizophrenia patients exhibit elevated prefrontal connectivity associated with longitudinal change. J Neurosci 35:267–286.

13. Cole MW, Anticevic A, Repovs G, Barch D (2011): Variable global dysconnectivity and individual differences in schizophrenia. Biol Psychiatry 70:43–50.

15. Driesen NR, McCarthy G, Bhagwagar Z, Bloch M, Calhoun V, D’Souza DC, et al. (2013): Relationship of resting brain hyperconnectivity and schizophrenia-like symptoms produced by the NMDA receptor antagonist ketamine in humans. Mol Psychiatry 18:1199–1204.

16. Driesen NR, McCarthy G, Bhagwagar Z, Bloch MH, Calhoun VD, D’Souza DC, et al. (2013): The impact of NMDA receptor blockade on human working memory-related prefrontal function and connectivity. Neuropsychopharmacology. 38:2613–2622.

17. Liang X, Zou Q, He Y, Yang Y (2013): Coupling of functional connectivity and regional cerebral blood flow reveals a physiological basis for network hubs of the human brain. Proc Natl Acad Sci U S A 110:1929–1934.

18. Cole MW, Pathak S, Schneider W (2010): Identifying the brain’s most globally connected regions. Neuroimage 49:3132–3148.

